# Herptile gut microbiomes: a natural system to study multi-kingdom interactions between filamentous fungi and bacteria

**DOI:** 10.1101/2023.08.23.554450

**Authors:** Lluvia Vargas-Gastélum, Alexander S. Romer, N. Reed Alexander, Marjan Ghotbi, Kylie C. Moe, Kerry L. McPhail, George F. Neuhaus, Leila Shadmani, Joseph W. Spatafora, Jason E. Stajich, Javier F. Tabima, Donald M. Walker

## Abstract

Reptiles and amphibians (herptiles) represent some of the more endangered and threatened species on the planet and numerous conservation strategies are being implemented with the goal of ensuring species recovery. Little is known, however, about the wild gut microbiome of herptiles and how it relates to the health of wild populations. Here we report results from both a broad survey of hosts and a more intensive sampling of hosts and geography of fungi and bacteria associated with herptile gut microbiomes. We demonstrate that bacterial communities sampled from frogs, lizards and salamanders are structured by the host higher level taxonomy and that the fungus *Basidiobolus* is a common and natural component of these wild gut microbiomes. Intensive sampling of multiple hosts across the ecoregions of Tennessee revealed that geography and host:geography interactions are strong predictors of distinct *Basidiobolus* OTUs present within a given host. Co-occurrence analyses of *Basidiobolus* and bacterial community diversity supports a correlation and interaction between *Basidiobolus* and bacteria, suggesting that *Basidiobolus* may play a role in structuring the bacterial community. We further the hypothesis that this interaction is advanced by unique specialized metabolism originating from horizontal gene transfer from bacteria to *Basidiobolus*, and demonstrate that *Basidiobolus* is capable of producing a diversity of specialized metabolites including small cyclic peptides.

**IMPORTANCE:** This work significantly advances our understanding of interactions in herptile microbiomes; the role that fungi play as a structural and functional member of herptile gut microbiomes; and the chemical functions that structure host:microbiome phenotypes. We also provide an important observational system of how the gut microbiome represents a unique environment that selects for novel metabolic functions through horizontal gene transfer between fungi and bacteria. Such studies are needed to better understand the complexity of gut microbiomes in nature and will inform conservation strategies for threatened species of herpetofauna.

## INTRODUCTION

Reptiles and amphibians (herptiles) are among the most threatened species on the planet. Approximately 20% of evaluated reptiles and 40% of amphibians are threatened with extinction (IUCN 2021 - International Union for Conservation of Nature and Natural Resources). Given the large threat to biodiversity, active conservation strategies are currently being utilized, including captive breeding programs, establishment of assurance populations and wildlife corridors, and translocation of individuals to enhance population genetics (1); however, the microbiome and how it relates to the health of wild populations has yet to be broadly incorporated into wildlife conservation. Pathogen induced dysbiosis (2), habitat degradation (3, 4) and climate change (5, 6) are all linked to alterations in the microbiome with potential for adverse consequences to the host organism. In depth knowledge of microbial interactions in the herptile gut microbiome is therefore necessary for conservation and management strategies, to facilitate understanding of perturbations to native microbiomes, as an increasing number of species are managed in captive breeding programs (7, 8).

The skin microbiome of amphibians has been the subject of considerable research due to chytridiomycosis caused by the fungus *Batrachochytrium dendrobatidis* (*Bd*), a skin pathogen of amphibians. The skin microbiome acts as a first line of defense against fungal pathogenicity (9) and its composition can be altered by factors such as microtopography, host life history, and environmental heterogeneity (10). Chytridiomycosis is known to alter the amphibian skin microbiome in field populations and laboratory experiments (11), correlates with distinct functional and metabolic properties (12), and results in poor community resilience following pathogen induced dysbiosis (13). Interestingly, it has been found that the skin microbiome of harlequin frogs (*Atelopus varius*) underwent rewilding when released into outdoor mesocosms from a captive breeding program (14).

While herptile-fungal interactions have been the subject of recent focus due to chytridiomycosis, the gut microbiome of herptiles remains understudied compared to other groups of animals. A Google Scholar search (January 18, 2023) for the term “Gut Microbiome” plus the following taxon generated the number of results reported: Amphibian= 5120, Reptile= 4420, Herptile= 13,900, Birds= 27,100, Mammals= 23,700, Mouse= 217,000, Human= 550,000. Furthermore, only a few studies have simultaneously focused on more than one domain of life, i.e., bacteria, in the herptile gut (e.g., (15–17). Although as many as 50 genera of fungi have been documented in the human gut, (18), fungi - mainly yeasts, make up a relatively small proportion of the human gut microbiome (19). Even less is known about fungi inhabiting the digestive systems of wildlife species. For example, the obligate anaerobic gut fungi (AGF), which are ubiquitously distributed among herbivorous ruminant animals and are essential to the digestion of lignocellulosic plant fiber, were only recently discovered in 1975 (20–22). Many herptiles are not herbivorous and instead feed on invertebrates like insects. Feeding strategy is predictive of the gut microbiome since herbivorous herptiles host different assemblages of gut microbiota compared to insectivores, including from the fungal genus *Basidiobolus* (16).

*Basidiobolus* is a filamentous, gut-inhabiting fungus isolated from the feces of a wide diversity of reptile and amphibian hosts (17, 23, 24). Resting spores of *Basidiobolus* are dispersed in fecal pellets, and upon defecation, germinate to produce hyphae and a diversity of spore types (Fig. 1). Hyphae grow and develop into a vegetative thallus (mycelium), and in many species produce sexually reproductive gametangia which fuse to form zygospores. Hyphae also give rise to conidiophores that produce apical, forcibly discharged asexual primary spores, blastoconidia, that germinate and give rise to a mycelium, or other spore types including capilloconidia. Capilloconidia possess adhesive tips which adhere to the exoskeletons of passing insects. These insects are eventually consumed by insectivorous hosts, reinoculating the host animal and completing the life cycle (Fig. 1). In addition to its herptile gut environment, *Basidiobolus* has been isolated from soil and leaf litter, and it is also known as an opportunistic pathogen of mammals (25) and frogs (26). Species of *Basidiobolus* have been documented from the gut of a wide diversity of metazoan hosts including anurans, bats, fishes, lizards, salamanders, snakes, turtles, and wallabies (23, 27–31).

**Fig 1.**
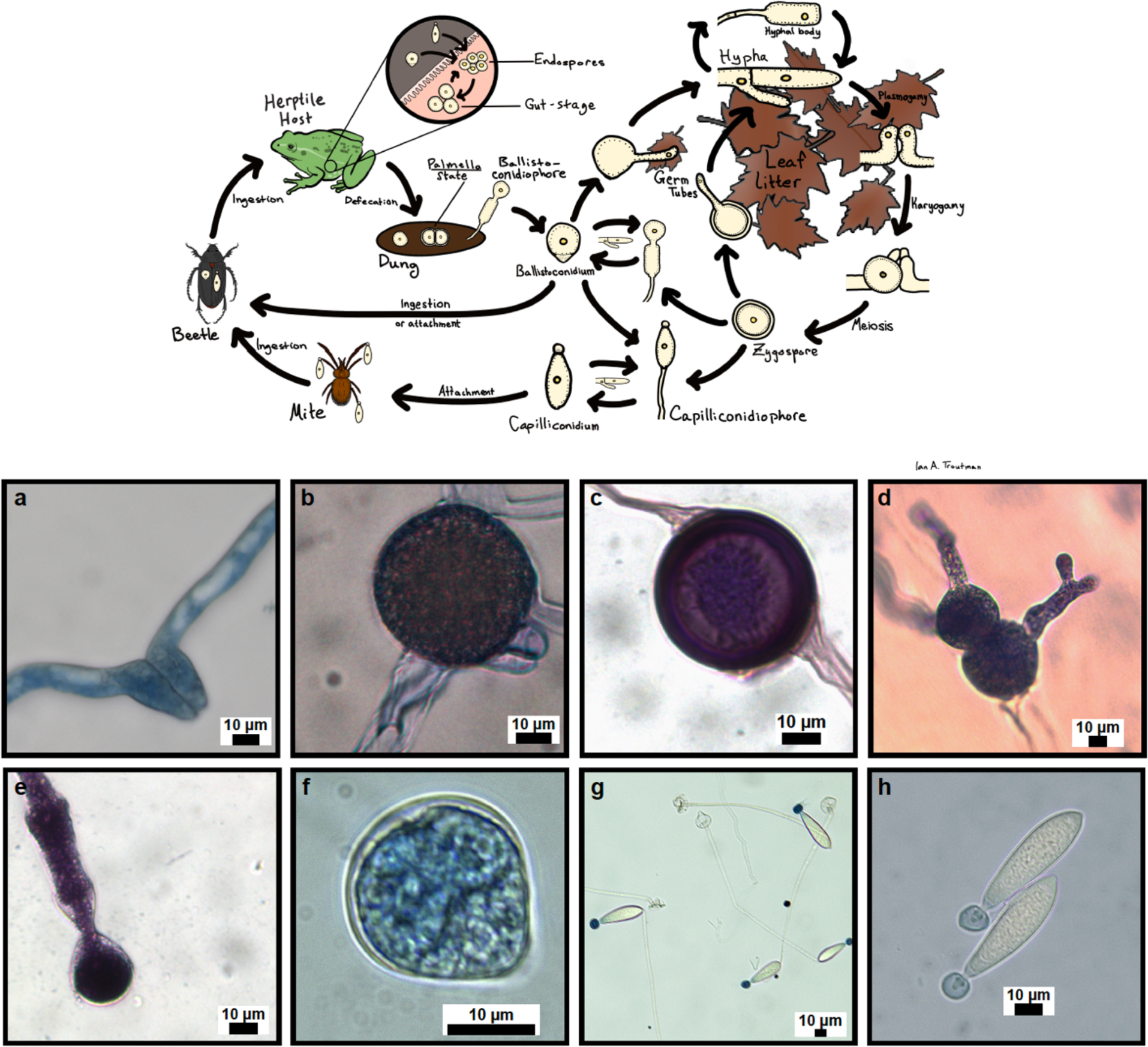
*Basidiobolus* life cycle and spore morphologies. Schematic of *Basidiobolus* life cycle showing major spore and vegetative stages and representation of spore morphologies: a) Compatible hyphae prior to zygospore formation, b) young zygospore, c) mature zygospore, d) zygospores germinanting, e) ballistoconidiophore, f) ballistoconidium, g) capilloconidiophores, h) capilloconidia.

This work is focused on understanding the diversity of bacteria and fungi in the gut of herptiles, interactions between *Basidiobolus* and other gut bacteria and fungi, genome signatures of HGT in *Basidiobolus,* and the suite of metabolites produced by species of *Basidiobolus*. We characterize the gut microbiome of 33 species of frogs, lizards and salamanders and show differences across both host and geographic space. We expanded the phylogenetic diversity of living *Basidiobolus* cultures and documented signatures of host and geographic preference and co-colonization of more than one putative species of *Basidiobolus* in the gut of herptile individuals. Network and indicator species analyses suggest correlations between *Basidiobolus* and other gut fungi and bacteria. Phylogenomic analysis of *Basidiobolus* indicated 2-5% of genes predicted to be of bacterial origins with enrichment of genes coding for specialized metabolism. Non-targeted LC-MS/MS and network analysis revealed peptidic metabolite signatures produced by cultures of *Basidiobolus* and in the gut of herptiles. We discuss herptile gut microbiomes in the context of fungal adaptations to the animal gut microbiome environment and microbial interactions between filamentous fungi and bacteria.

## RESULTS AND DISCUSSION

### Herptile microbiomes are characterized by unique bacterial and filamentous fungal communities

The rarefied dataset consisted of 133 herptiles (herptilesTable S1) from eight different states (Arizona, Ohio, North Carolina, Tennessee, Georgia, Alabama, Arkansas, and Louisiana). A total of nine samples were removed from the 16S rRNA and ITS rDNA datasets due to quality control filtering. After quality control, DADA2 processing, decontamination and rarefaction, a total of 9,196,775 fungal ITS rDNA sequences and 6,110,012 bacterial 16S rRNA sequences were retained. These sequences resulted in a total of 5,562 fungal ASVs and 9,343 bacterial ASVs. Nine fungal and 35 bacterial phyla were identified. The most abundant phyla were Ascomycota, Basidiobolomycota, and Basidiomycota for fungi, and Bacteroidota, Firmicutes, and Proteobacteria for bacteria (Fig. 2).

**Fig 2.**
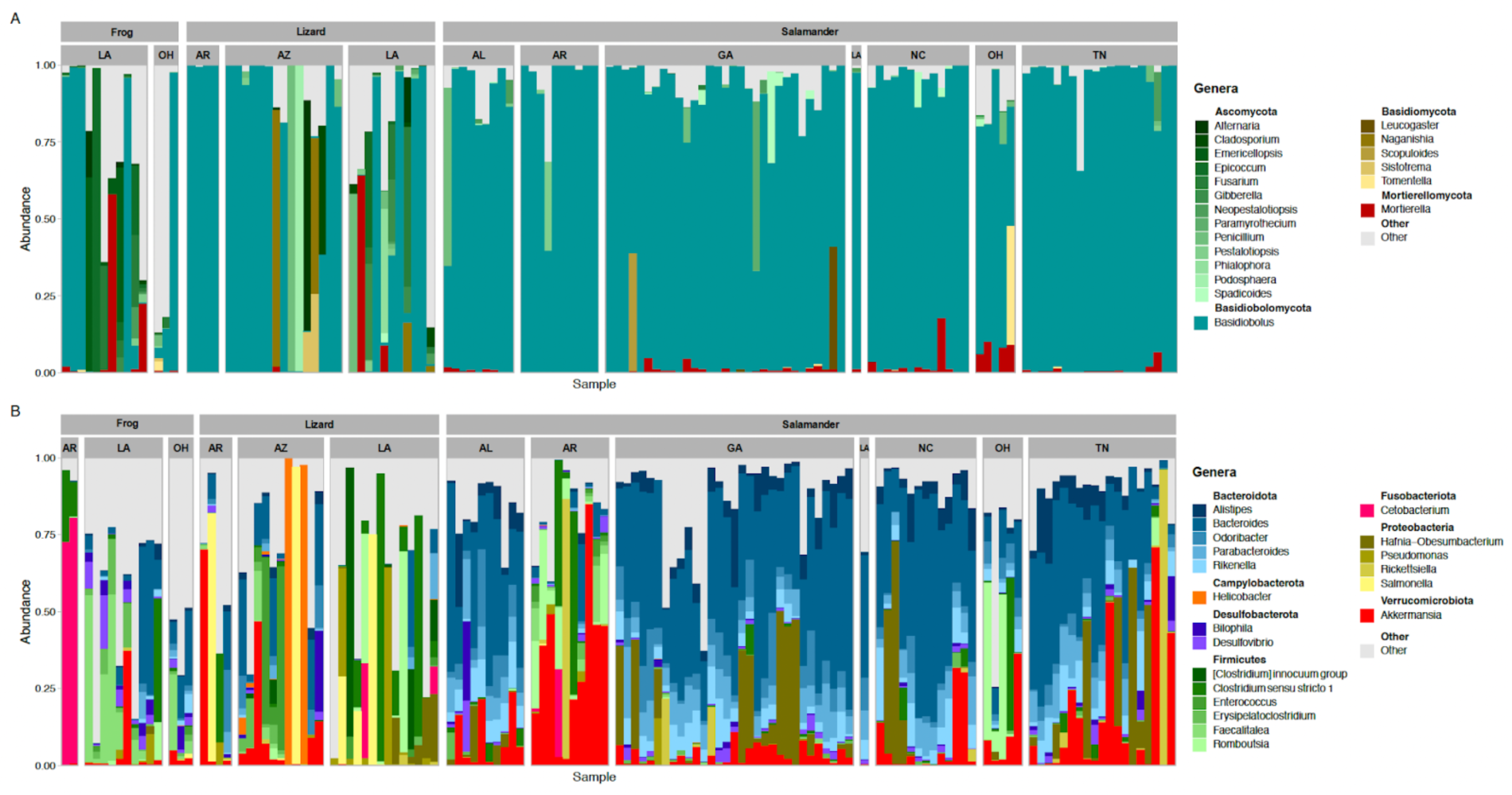
Taxonomic composition of the twenty most abundant genera of the herptile gut microbiome. Fungi (A) and bacteria (B) found in the gut of frogs, lizards, and salamanders from the different sampled geographic locations. Alabama (AL), Arkansas (AR), Arizona (AZ), Georgia (GA), Louisiana (LA), North Carolina (NC), Ohio (OH), Tennessee (TN).

Differences were observed when comparing relative abundance data between hosts for both bacteria and fungi (Fig. 2). For bacteria, the most obvious difference was the differential representation of Bacteriodota and Firmicutes as a function of host grouping of frogs, lizards, and salamanders. Firmicutes represented 64.7% of abundance in frogs; Firmicutes represented 39.7% and Proteobacteria 39% of the abundance in lizards; and Bacteroidota accounted for 40.1% of abundance in salamanders. Relative abundances of salamander gut bacteria were mostly similar, except for Arkansas and some Ohio sites, where they were dominated by Firmicutes and *Akkermansia* (Fig. 2B).

For fungi, *Basidiobolus* represented the most abundant group in salamanders and lizards with values of 84.4% and 52.8%, respectively. In frogs, the Ascomycota was the most abundant group (44.2%) followed by *Basidiobolu*s (29%). *Basidiobolus* dominated the fungal composition in the majority of salamander samples, ranging from >60% to 99% of abundance, and little variation in fungal communities across different geographic locations. Frog and lizard samples were more variable across geographic localities, as the majority of samples were dominated by *Basidiobolus,* or other members of Ascomycota (*Alternaria*, *Emericellopsis* and *Epicoccum, Penicillium*) and Basidiomycota (*Naganishia* sp.; Fig. 2A). Several other species of fungi have been documented in the gut of herptiles including *Aspergillus fumigatus*, *Geotrichum candidum*, *Trichosporon* sp., and *Candida parapsilosis* (23). In a metabarcoding and high throughput ITS rDNA sequencing study (16) it was found that *B. ranarum* and *B. magnus* dominated the core fecal mycobiome of *Sceloporus grammicus* lizards. They also documented *Aureobasidium microstictum*, *Hyphopichia burtonii*, *Penicillium thomii*, *Talaromyces duclauxii*, and *Tetraspisispora fleetii* as members of the lizard mycobiome.

The PERMANOVA assumption of homogeneity of variance was violated when comparing multivariate dispersion of host groups for both bacteria (betadisper; F_2, 132_ = 8.205, p=0.002) and fungi (betadisper; F_2, 132_ = 10.026, p=0.001). However, PERMANOVA is robust to this violation since the group with the largest sample size (salamanders) showed the most variance in multivariate dispersion for both bacteria and fungi (32). A significant effect of host (F_2, 132_ = 7.915, R^2^ = 0.096, p = 0.001), geography (F_7, 132_ = 3.258, R^2^ = 0.138, p = 0.001) and the interaction term (F_3, 132_ = 2.014, R^2^ = 0.037, p = 0.001) was observed for average bacterial assemblages. For fungi, a significant effect of host (F_2, 132_ = 9.989, R^2^ = 0.113, p = 0.003), geography (F_7, 132_ = 4.432, R^2^ = 0.175, p = 0.004) and the interaction term (F_3, 132_ = 2.406, R^2^ = 0.027, p = 0.001) was observed for average assemblages. These patterns were substantiated by the PCoA plots for both bacteria and fungi (Fig. 3) and reinforced that herptile gut microbiomes are shaped by host (e.g., (33), geography (e.g., (17), and their interactions.

**Fig. 3.**
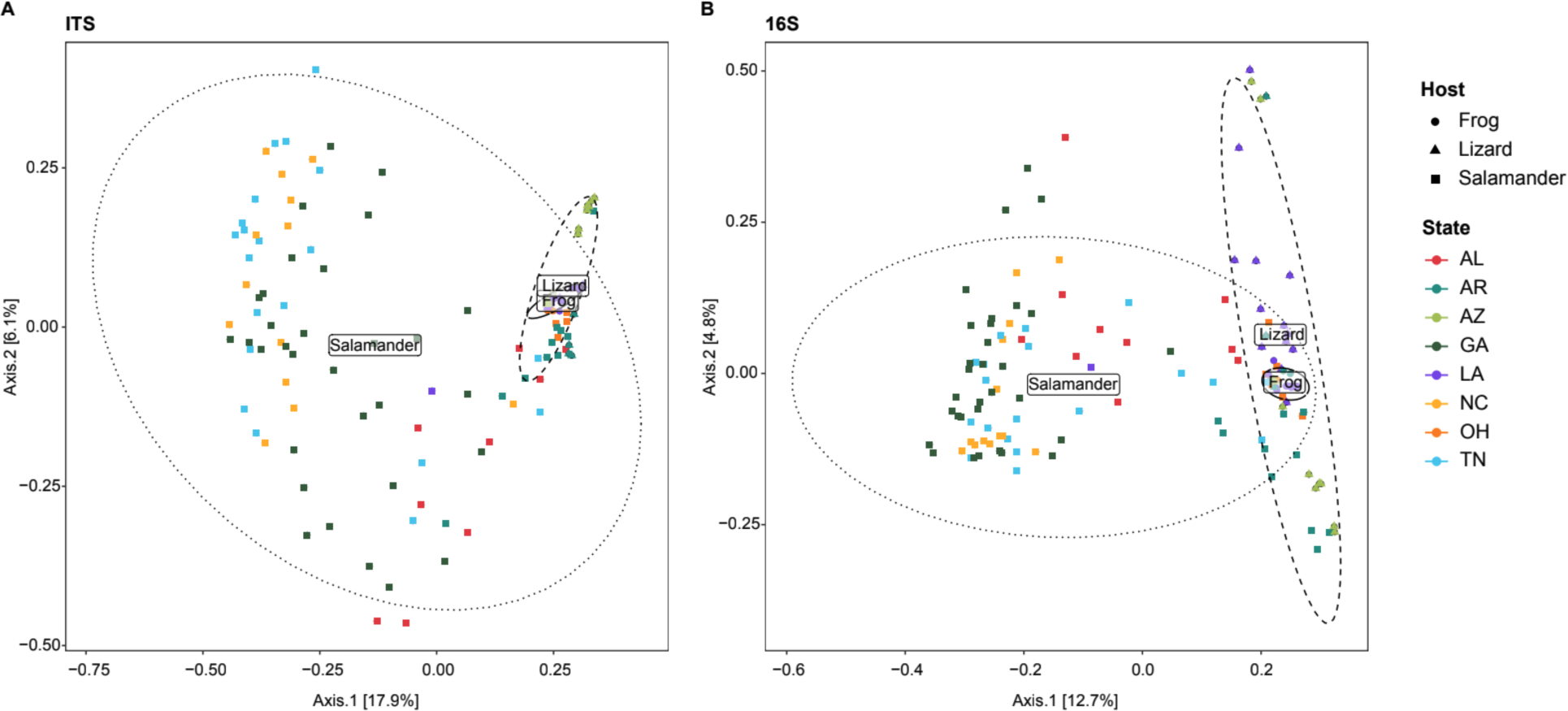
PCoA of fungal and bacterial communities based on Bray-Curtis dissimilarity. Host and geographic diversity of gut fungi (A) and bacterial (B) assemblages. Alabama (AL), Arkansas (AR), Arizona (AZ), Georgia (GA), Louisiana (LA), North Carolina (NC), Ohio (OH), Tennessee (TN).

### The herptile gut microbiomes harbor phylogenetically diverse *Basidiobolus* OTUs

A total of 336 *Basidiobolus* ITS rDNA sequences were aligned and analyzed phylogenetically (phylogeneticallyFig. S1). Fifty-two corresponded to reference sequences downloaded from NCBI, and the remaining 284 corresponded to living isolates obtained in this project. Phylogenetic analysis revealed that *Basidiobolus* isolates represent nine well supported clades (>87%, outlier of 54%). The tree is composed of two principal clades with strong bootstrap values (85%): one consisting of a group of 26 sequences archived on NCBI GenBank and three new *Basidiobolus* isolates collected in this project from a single frog (*Lithobates clamitans*) individual, and a second clade represented by 26 reference sequences and 281 study sequences (Fig. 4), many of which were associated with one host species, with some showing geographic specificity at the EPA Ecoregion IV level (sequencese colored bars, Fig. S1).

**Fig. 4.**
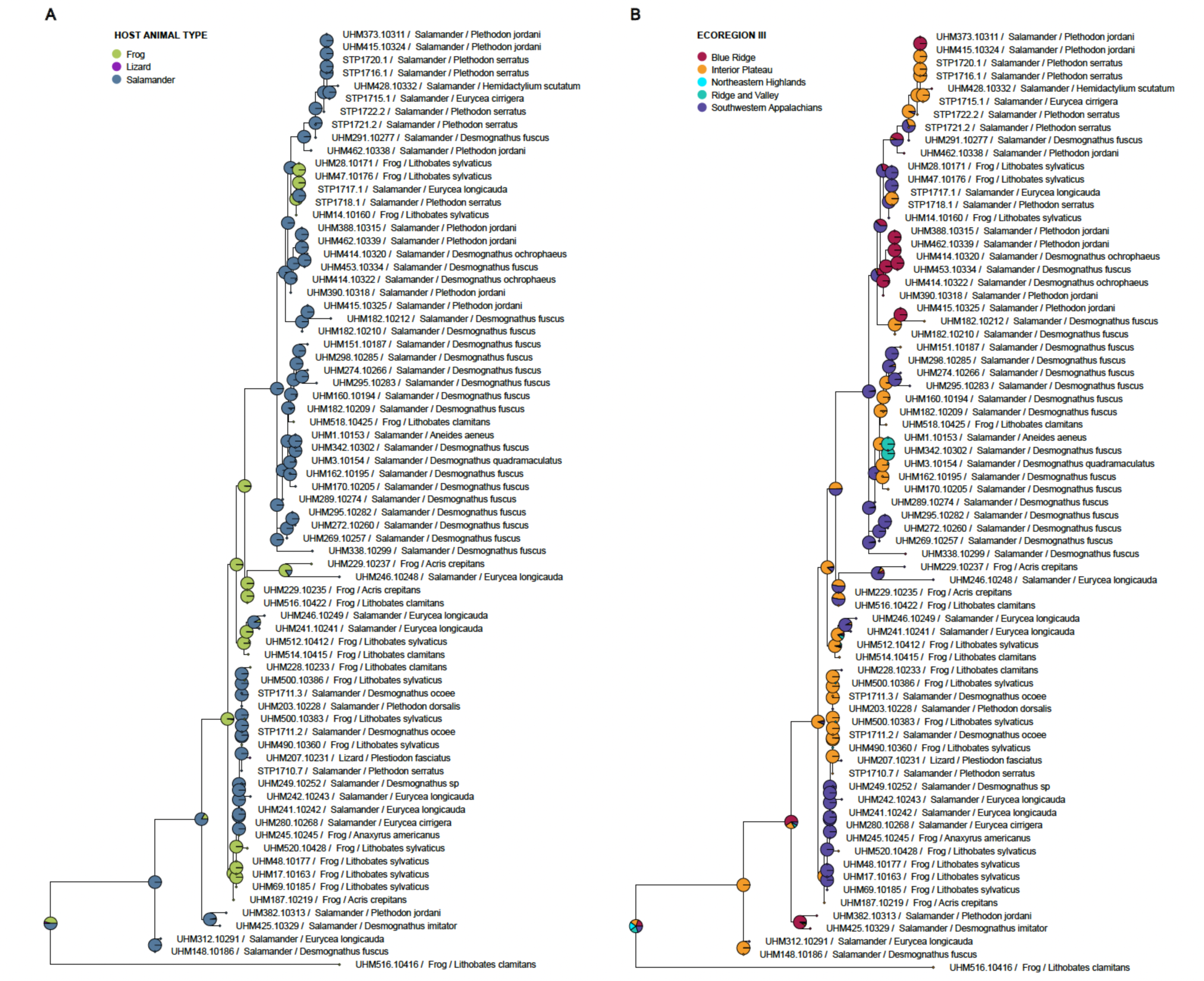
Ancestral Character State Reconstruction. Analysis testing host groups (A) and ecoregion III (B) classifications. Pie charts are indicative of the relative likelihoods of each node being in a particular state.

A single ITS sequence variant of *Basidiobolus* was collected from most herptile individuals using Sanger sequencing (n=274 of 284 individuals). But ten individual amphibian hosts were co-colonized by genetically different isolates of *Basidiobolus* (italicFig. S1), demonstrating the ability of the herptile gut microbiome to harbor multiple *Basidiobolus* OTUs. Similar patterns of host-fungal specificity have been observed in the gut mycobiome of termites (34) and Slimy Salamanders (17).

In order to compare current and past work (17), we used comparisons of operational taxonomic units (OTUs), amplicon sequence variants (ASVs) and phylogenetic analysis of Sanger sequence data. There are only 10 described species of *Basidiobolus*, but ITS rDNA data from amplicon metabarcoding studies that sampled herptile fecal samples are consistent with there being substantial undescribed biodiversity (biodiversityFig. S1). However, allelic variation in the *Basidiobolus* ITS rDNA marker complicates this interpretation. For example, from 59 Slimy Salamander fecal samples Walker et al. (2020) found 485 *Basidiobolus* OTUs clustered at 97% similarity, and only two names could be provisionally linked to five of the OTUs with species epithets. Furthermore, only four species of *Basidiobolus* (*B. heterosporus, B. magnus, B. microsporus, B. ranarum*) are found in the UNITE v.9.0 reference database (34). It is unlikely that all 485 OTUs were representatives of phylogenetic species, as we have found 4–14 ITS rDNA amplicon sequence variants (ASVs) in the DNA of six living cultures of *Basidiobolus* (Table 1), and *B. meristosporus* is estimated to have more than a thousand ITS rDNA copy variants in its genome (35). However, PCR and Sanger sequencing of genomic DNA isolated from cultures produced a single ITS product, which mapped to the dominant ASV detected in cultures via Illumina sequencing of amplicons (Fig. S2).

**Table 1.** Allelic diversity of ITS rDNA marker gene in living isolates of *Basidiobolus*. Data were generated using 2 x 250 bp paired end sequencing on an Illumina MiSeq and analyzed in DADA2.

| Isolate ID | Fungal ID | Host ID | Total number of<br>*Basidiobolus*<br>ASVs | Abundant<br>*Basidiobolus* ASVs<br>(n>5 reads) |
| --- | --- | --- | --- | --- |
| STP1710.7 | *Basidiobolus* sp. | *Eurycea longicauda*<br>(Long-tailed Salamander) | 13 | 12 |
| STP1718.1 | *Basidiobolus* sp. | *Plethodon serratus*<br>(Southern Red-backed<br>Salamander) | 4 | 4 |
| STP1717.1 | *Basidiobolus* sp. | *Plethodon serratus*<br>(Southern Red-backed<br>Salamander) | 6 | 4 |
| STP1715.2 | *Basidiobolus* sp. | *Eurycea cirrigera*<br>(Southern two-lined<br>Salamander) | 14 | 12 |
| UHM1.3285 | *Basidiobolus* sp. | *Aneides aeneus*<br>(Green Salamander) | 6 | 6 |
| UHM3.3284 | *Basidiobolus* sp. | *Desmognathus*<br>*quadramaculatus*<br>(Blackbelly Salamander) | 8 | 6 |

The Ancestral Character State Reconstruction Analysis (ACSR) of frog, lizard and salamander resolved a mixing of hosts across the ITS phylogeny (Fig. 4A). Salamanders were resolved as the dominant ancestral host of *Basidiobolus*, which may indicate an evolutionarily and/or ecologically meaningful interaction between host life history and the *Basidiobolus* life cycle, an interpretation consistent with a higher frequency of *Basidiobolus* detected in salamander fecal samples versus frogs and lizards (Fig. 2A). The ACSR analyses of geography also indicated a mixing of ecoregions suggesting that some OTUs, or closely related OTUs, are distributed across multiple ecoregions (e.g., Interior Plateau and Southwestern Appalachians) while others are more restricted in their distributions (e.g., Blue Ridge and Ridge and Valley) (Fig. 4B).

Patristic distances from the ITS tree were used to test the differential effect of host and geography, and their interaction, across the *Basidiobolus* isolates sampled. Multivariate dispersion was not significantly different among hosts groups (betadisper; F_2, 70_ = 0.7065, p=0.394), host genera (betadisper; F_8, 64_ = 1.8513, p=0.102), or ecoregion III (betadisper; F_4, 68_ = 2.2075, p=0.126). There was a significant effect on *Basidiobolus* genetic variation based on host group (frogs, lizards, salamanders; F_2, 72_ = 3.6322, R^2^ = 0.0659, p=0.016), ecoregion III (F_2, 72_ = 6.3010, R^2^ = 0.2286, p=0.002) and the interaction term (F_2, 72_ = 6.8714, R^2^ = 0.1247, p=0.020). A finer scale assessment of host genus and the interaction term revealed only marginally significant effects (Genus: F_8, 72_ = 2.2488, R^2^ = 0.1529, p=0.086; Genus x Ecoregion III: F_8, 72_ = 3.2024, R^2^ = 0.1633, p=0.070) on *Basidiobolus* genetic distance. While these results are consistent with geographic effect and host:ecoregion interaction being more significant explanatory variables than host association, additional collections of *Basidiobolus* isolates from other hosts (e.g., lizards and frogs) will allow for more robust testing of hypotheses related to host association.

### *Basidiobolus* OTUs and bacterial gut communities are co-structured

Mostly research on herptile gut microbiomes has been descriptive in nature, focused entirely on bacteria, and with limited inference into microbial interactions (e.g. (36, 37). Multiple factors (e.g., diet, host taxonomy, disease) are hypothesized to modulate gut bacterial assembly though their relative contribution to this process has not been elucidated (38). For example, a study on Ornamented Pygmy Frogs (*Microhyla fissipes*) revealed the complex remodeling of gut bacteria during metamorphosis and found a possible coevolution between gut microbial groups and host dietary shifts (39). Large-scale restructuring in the Burmese Python (*Python bivittatus*) gut microbiome was observed to correspond with withphysiological changes of the host gut during snake feeding and fasting (40). The bacterial component of the herptile gut microbiome has been linked with diet (41), parasitic worm load (42), specific digestive system organs (33), and may exhibit metagenomic plasticity (43). A large scale characterization of the gut bacterial microbiome of Mammalia, Aves, Reptilia, Amphibia and Actinopterygii showed that diet selects for specific functional guilds while host evolutionary history selects for prevalence of particular OTUs (38). Although we have a rudimentary understanding of herptile gut bacteria, no study to date has attempted to characterize the structure, function, and interactions between more than one domain of life composing natural herptile gut microbiomes.

Walker et al., (2020) determined that species in the genus *Basidiobolus* averaged 60% (minimum 8.1%; maximum 97.8%) of the relative abundance of all gut fungi among individuals in the Slimy Salamander species complex. We demonstrate here that bacterial OTUs in the Slimy Salamander gut microbiome were correlated with the relative abundance of *Basidiobolus* OTUs (Fig. 5A). *Basidiobolus* OTUs classified as belonging to the same species are predicted by similar bacterial OTUs and clustered nearer to one another (Fig. 5A). Several unidentified species of *Basidiobolus* were also determined to correlate with at least 10 classes of bacteria including the Verrucomicrobiae, Bacteroidia and Clostridia (Fig. 5B). Mean indicator power values (44) of each bacterial class were used to determine if there was taxonomic variation in the ability of bacterial OTUs to predict *Basidiobolus* occurrence in Slimy Salamander gut microbiomes. A single OTU in the class Spirochaetia was the strongest indicator (IP = 0.446) of *Basidiobolus* (Fig. 5B). Numerous OTUs in the Bacteroidia (n=36) and the Clostridia (n=19) were weaker, but more abundant indicators of *Basidiobolus*. We examined whether the occurrence of certain *Basidiobolus* OTUs were correlated with the assemblage of bacterial OTUs using total indicator power (TIP; Fig. 5C). Results suggest that *Basidiobolus* OTUs do not interact equally with bacterial assemblages in the Slimy Salamander gut microbiome. *Basidiobolus ranarum* OTUs had higher TIP values than *B. heterosporus* (Fig. 5C) suggesting that different species of *Basidiobolus* display variability in the strength of their interactions with gut bacteria.

**Fig. 5.**
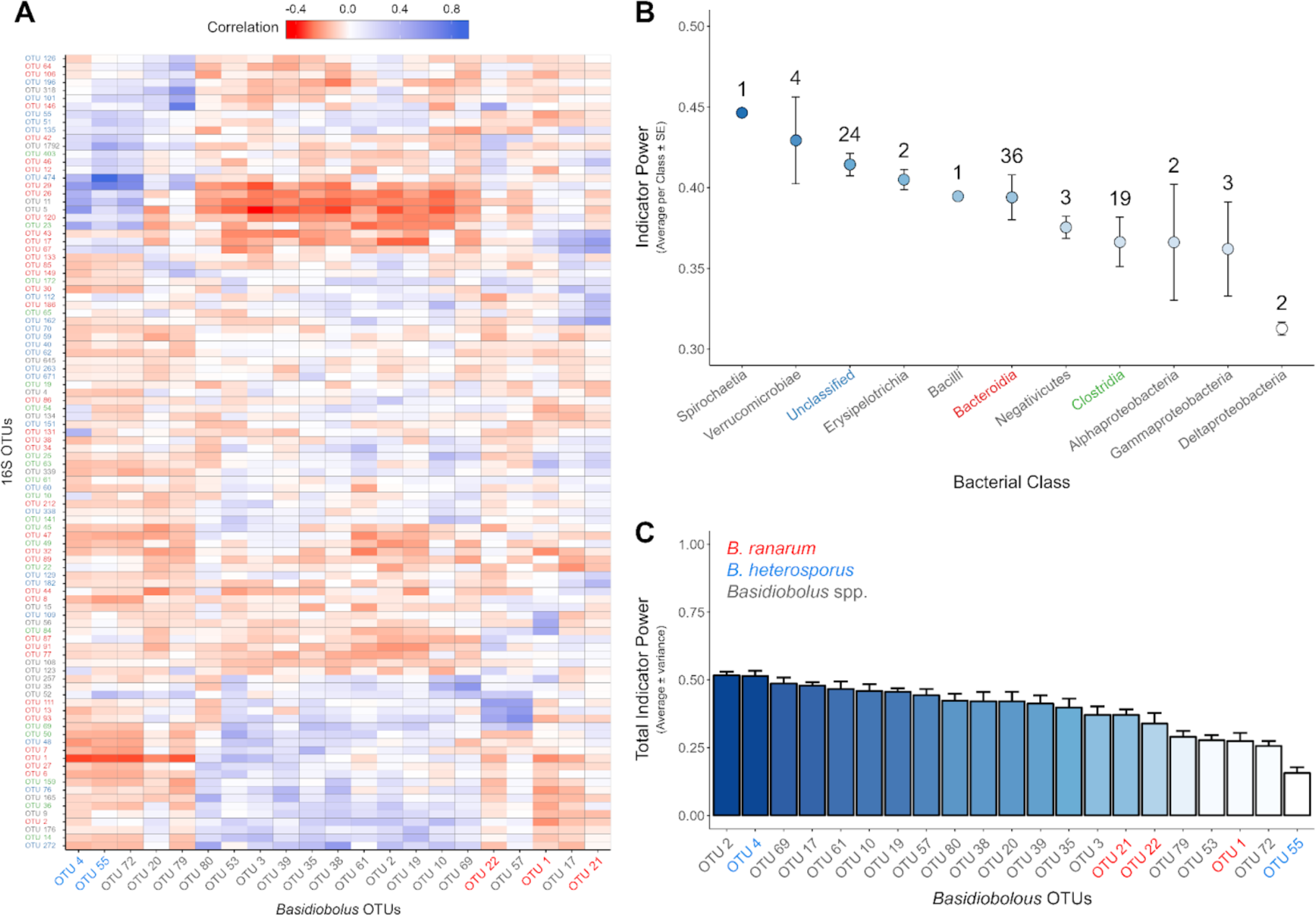
Indicator power analysis. (A) Heatmap of the Pearson correlation coefficients between the most abundant bacterial OTUs (n = 100) and abundant *Basidiobolus* OTUs (reads ≥ 10; n = 21 OTUs). Cooler heatmap colors indicate that a bacterial OTU is positively correlated with a *Basidiobolus* OTU. Conversely, warmer heatmap colors indicate that a bacterial OTU is negatively correlated with a *Basidiobolus* OTU. Hierarchical clustering was performed on the correlation matrix using the complete linkage algorithm. The clustering methodology was used to arrange the *Basidiobolus* OTUs on the x-axis and bacterial OTUs on the y-axis. (B) Using the 100 most abundant bacterial OTUs, mean indicator power was calculated for each bacterial class. This value represents the ability of an OTU to predict the occurrence of all *Basidiobolus* OTUs present in the dataset. Thus, OTUs from bacterial classes with high mean indicator power are strong predictors of the presence or absence of *Basidiobolus*. The number of OTUs from each bacterial class are annotated above each point. (C) Total indicator power, a measure of the ability of one *Basidiobolus* OTU to predict a complete assemblage of bacterial OTUs, was calculated for each *Basidiobolus* OTU. Thus, *Basidiobolus* OTUs with low total indicator power values (e.g. OTU55) are not well correlated with the overall structure of the herptile gut microbiome. *Basidiobolus* OTU labels classified using the UNITE as belonging to the same species are assigned distinct colors: red - *B. ranarum*, blue - B. *heterosporus*, gray - *Basidiobolus* spp.

We found the overall structures of the co-occurrence networks to be different between host animals (Fig. 6). The edge density (D) represents how dense the network is in terms of edge connectivity and significance of associations. Frog gut microbiomes had the lowest edge density (D=0.0207) compared to salamanders (D=0.0257) and lizards (D=0.0261). The transitivity (T), or clustering coefficient was the highest in frogs (T=0.1820) with their network containing 15 modules, followed by lizards (T=0.1704; 13 modules) and salamanders (T=0.1662; 10 modules). Higher clustering coefficient denotes presence of communities or groups of nodes that are densely connected internally and forming modules. Modules in microbial co-occurrence networks may infer ecological processes within microbiomes that influence microbial community structure, such as niche filtering and habitat preference, between specific groups of microbes (Lima-Mendez et al., 2015). Modules may also reflect the presence of functional and metabolic interactions between microbes that may be syntrophically coupled (45). Intensity of the within and between module connections is displayed through modularity of each network. The highest modularity was detected in lizards (0.781) compared to frogs (0.625) and salamanders (0.529). This may indicate the presence of stronger metabolic interactions between microbes shaping functional modules within the gut of lizards compared to frogs and salamanders. The frequency of positive interactions between pairs of microbes was high in all three hosts with only 8% of negative interactions (red vectors, Fig. 6A) in frogs, 0.46 % in salamanders (Fig. 6C) and 0% in lizards (Fig. 6B). The mostly positive correlations in networks may reflect functional guilds of microorganisms with cooperation due to similar or complementary functions, whereas, negative correlations may represent competition or parasitism.

**Fig 6.**
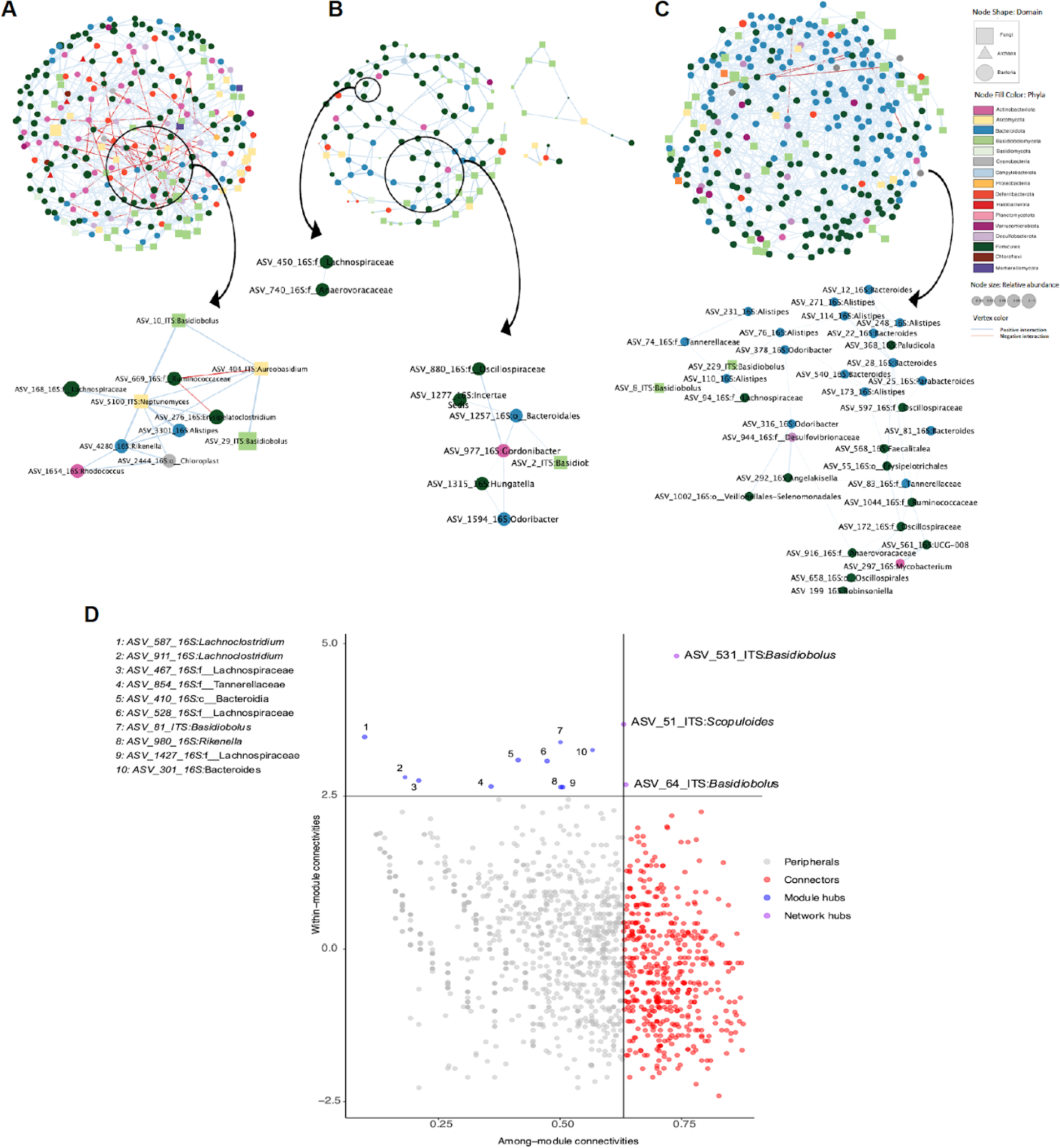
Network analysis of archaeal, bacterial and fungal microbiomes. Networks from (A) frogs, (B) lizards, and (C) salamanders. Each node represents an ASV and is shaped according to the taxonomic domain. Edge color denotes a positive (blue) or negative (red) interaction between two connected ASVs. The ASVs with main interactions chosen based on highest degree and betweenness are shown enlarged on the plot. (D) Scatter plot showing distribution of archaeal, bacterial and fungal ASVs according to their within-module and among-module connectivity. Each dot represents an ASV in the complete dataset of all herptile hosts. The four panels show the role distribution of selected groups of microbes represented by four different colors. ASVs representing network hubs are indicated by numbers on the plot and listed in the upper left panel (1–10).

Cross domain microbial interactions differed between animal hosts (Fig. 6). In frogs we found the highest inter domain diversity (Archaea, Bacteria and Fungi) of interactive microbes. Archaeal ASVs were not detected in lizards and salamanders. Subnetworks were created representing the connectivity of a node with other nodes in a network (Fig. 6D; top 30% degree values). In frogs, an ASV of *Aureobasidium* (ASV-404-ITS; Ascomycota) had the highest betweenness and centrality, demonstrating a dominant role in the network. *Aureobasidium* showed negative interactions with ASV669-16S from the family Ruminococcaceae (phylum Bacillota) and positive interactions with two *Basidiobolus* ASVs (ASV-10-ITS, ASV-29-ITS), *Neptunomyces* (ASV-5100-ITS; phylum Ascomycota) and *Alistipes* (ASV-3301-16S; phylum Bacteroidetes). Two separate subnetworks were highlighted for lizards. They included positive interactions between bacteria from the phylum Bacillota (ASV-450-16S: *Lachnospiracaea*, ASV-740-16S: *Anaerovoracaceae* dual node cluster and ASV-880-16S, ASV-1277-16S, ASV-1315-16S: *Hungatella* main cluster), the Bacteroidetes (ASV-1257-16S, ASV-1597-16S: *Odoribacter*), the Actinomycetota (ASV-977-16S: *Gordonibacter*) and an ASV of *Basidiobolus* (ASV-2-ITS). Likewise, salamander subnetworks consisted of ASVs in the phyla Bacillota, Bacteroidetes, and Actinobacteriota. The only fungal ASVs selected with a dominant role occurred in the genus *Basidiobolus* (ASV-8-ITS, ASV-229-ITS). Overall, in all three host groups, fungal and bacterial co-occurrence networks detected strong and consistent interactions between nodes annotated as *Basidiobolus* and bacterial nodes belonging to Bacteroidota and Firmicutes.

To identify key functional groups in herptile gut microbiomes, nodes were classified into four categories of peripherals, connectors, module hubs and network hubs (46, 47). Peripheral ASVs can be interpreted as specialists, whereas, module hubs and connectors are generalists, and network hubs super-generalists (47). Connectors, generalists, and super-generalists are considered to be keystone microorganisms playing a critical role in network structure (48). Two ASVs of *Basidiobolus* and one *Scopuloides* (crust fungi belonging to Basidiomycota) were identified as network hubs (Fig. 6D). Nine ASVs belonging to Bacteroidetes and Firmicutes and one ASV of *Basidiobolus* comprised module hubs, denoting key roles of these microbes in the structure and stability of herptile gut microbiomes.

### Patterns of fungal-bacterial co-occurrences highlight the importance of *Basidiobolus*

Deciphering the interactions among fungal-bacterial species within their complex and contiguous communities is a pivotal goal of microbial ecology. Network-based analytical approaches have proven to be a powerful tool to study interactions through co-occurrences within multi-domain complex microbial systems (49). Co-occurrence analysis was performed at the ASV level with taxonomic identification to genus level. In order to identify more stable bacterial and/or fungal interactions for each host animal group, only ASVs present in more than 20% of samples were included (Fig. 6). The overall structures of the networks differed between frogs, lizards and salamanders. The edge density (D) of the network, which represents how dense the network is in terms of edge connectivity or number of cooccurrences, was the lowest in frogs (0.0207) compared to salamanders (0.0257) and lizards (0.0261). However, the transitivity (T), or clustering coefficient was the highest in frogs (0.1820), followed by lizards (0.1704) and salamanders (0.1662). Higher clustering coefficiency denotes presence of communities or groups of nodes that are densely connected internally and may reflect the presence of functional modules (50). The frequency of positive interactions between pairs of microbes was higher in all three herptile host gut microbiomes. The highest proportion of negative edges was seen in frogs (8%), followed by salamanders (0.46 %) and lizards (0%) (x0025Table S3).

### Horizontal gene transfer and its connection to specialized metabolism in *Basidiobolus*

The best documented case of HGT in fungi involves the anaerobic gut fungi (AGF), which are zoosporic fungi that reside in the GI tract of ruminant animals and are an essential biological component of the ruminant bioreactor (50). AGF genomes have a documented HGT rate of 2.0-3.5%, allowing AGF to expand substrate utilization range, diversify pathways for electron disposal, acquire novel secondary metabolism, and facilitate adaptation to the anaerobic environment (51). *Basidiobolus* is the only known genus of fungi that is specialized to the herptile gut and represents an independent origin of gut fungi as compared to the AGF (52). Phylogenomic analyses of three draft *Basidiobolus* genomes reveal a similar magnitude of HGT as AGF, however, with 4-5% of genes predicted to have bacterial origins, with the largest sources being Actinobacteria, Firmicutes, and Proteobacteria (53). The most pronounced signal of HGT is in secondary or specialized metabolism for nonribosomal peptide synthetases, which are known to function in immunoregulation, quorum sensing, iron metabolism and siderophore activity (NRPS; NRPSFig. S3). It seems plausible that the herptile gut environment promotes HGT from bacteria to fungi under the selection pressure of acquisition of novel metabolism necessary to adapt to herptile microbiomes.

To test for production of metabolites consistent with NRPS biosynthesis, we cultured nine *Basidiobolus* isolates from six salamander individuals, as well as *B. meristosporus* CBS 931.73, in parallel on potato dextrose agar (PDA) for LC-MS/MS profiling. Each *Basidiobolus* culture plate produced 150 to 500 mg of dried mycelium and resulted in an average of 4.2 mg of extract (average of 18.7 mg of extract per gram dried mycelium). Data processing in MzMine resulted in selection of 331 mass features (m/z & retention time) with associated quantification (area under analyte chromatographic peak/area under internal standard chromatographic peak) across all samples. Feature-based molecular networking (54) of the resulting MS/MS spectra using the GNPS online platform (55) yielded nodes and subnetworks containing mass features from all 10 *Basidiobolus* cultures (figFig. 7). The GNPS feature based molecular network assigned 613 edges between the 331 nodes and spectral library annotations for 21 of those nodes. General chemical class assignments for larger sub-networks were determined by comparison of GNPS annotations (when applicable) with both SIRIUS and CANOPUS outputs for many of the networked nodes. While the identity of each node cannot be determined at this level of analysis, a general overview shows one large subnetwork containing fatty acids, three sub-networks of phosphocholines, two cyclic peptide sub-networks, one containing steroids, and one with sphingolipids. Excluding the steroid subnetwork (pink), primarily derived from STP1710.1, the remaining chemical classes are generally shared between all *Basidiobolus* isolates. For example, both cyclic peptide subnetworks contain nodes representing mass features present in more than one of the isolates. Nodes with contributions from different *Basidiobolus* isolates in the two cyclic peptide subnetworks are consistent with a conservation of biosynthetic potential for these specialized metabolites at the genus level. Notably, in both cyclic peptide subnetworks, *Basidiobolus* isolates from the same animal share the same structurally related mass features (e.g., STP1718.2 and STP1718.4) that are also linked to other structurally related mass features (nodes) found in extracts of other *Basidiobolus* isolates. In some cases, *Basidiobolus* isolates from different animals of the same species share some cyclic peptide mass features, e.g., STP1717.1 and STP1718.2 from two different Southern Red-backed Salamander (*Plethodon serratus*) individuals. The conservation of multiple specific cyclic peptide mass features between *Basidiobolus* isolates from different salamander species is also striking. For example, *Basidiobolus* isolate STP1710.7, from a Long-tailed Salamander (*Eurycea longicauda*), and STP1711.2 and STP1711.3 isolates from the same Ocoee Salamander (*Desmognathus ocoee*) share multiple nodes in both cyclic peptide subnetworks.

**Fig 7.**
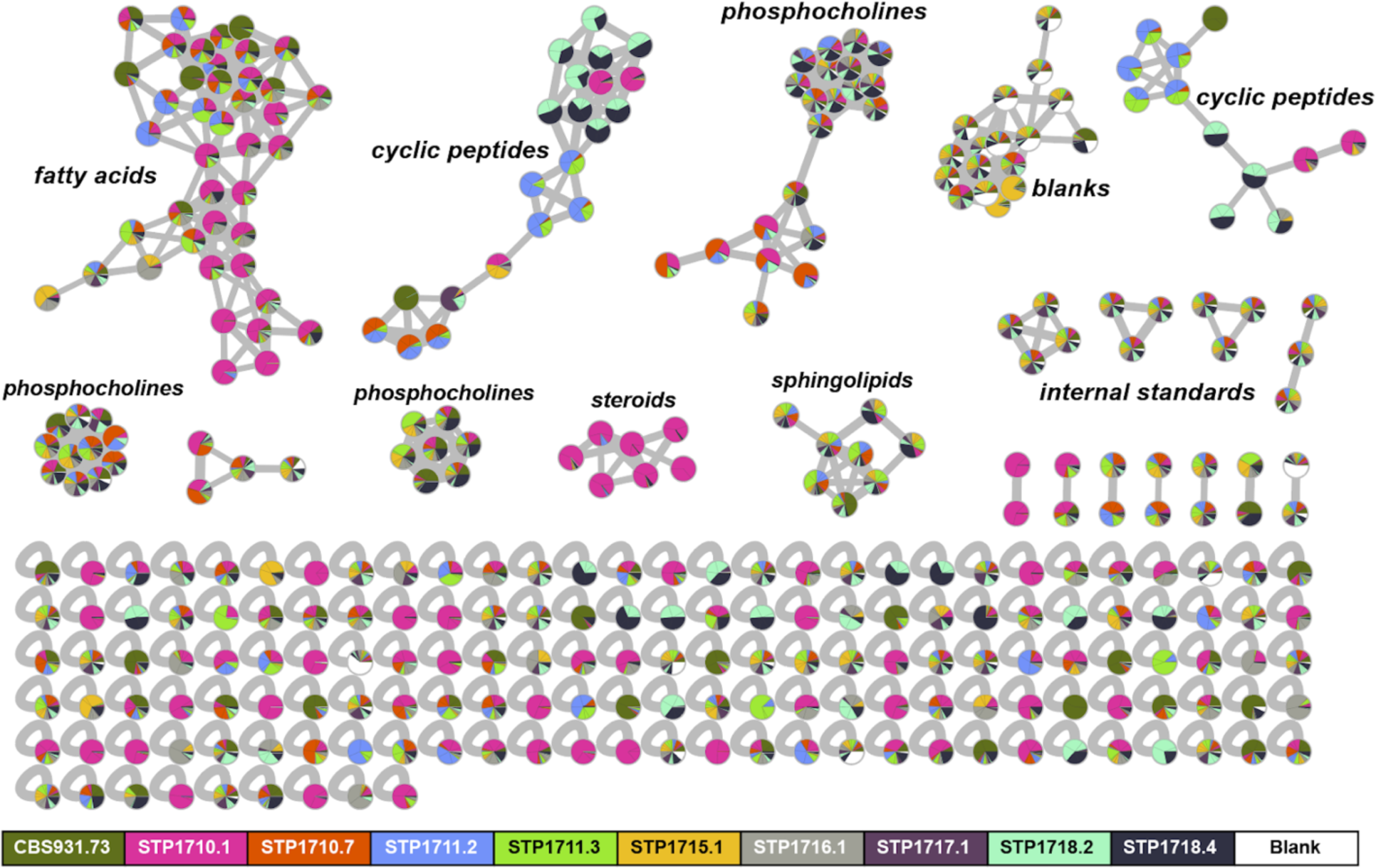
GNPS feature-based molecular network of untargeted LC-MS/MS data for extracts of 10 different *Basidiobolus* isolates cultured on PDA. Mass features are represented as nodes and are colored according to source isolate, with pie charts representing mass features shared between *Basidiobolus* isolates. Edges connect mass features (nodes) with similar MS/MS spectra, defined as a cosine similarity score > 0.7, which indicates structural relatedness. *Basidiobolus* isolates derive from feces of a gecko (*B. meristosporus* CBS 931.73) and salamanders *Eurycea longicauda* (STP1710.1, STP1710.7, one animal), *Desmognathus ocoee* (STP1711.2, STP1711.3, one animal), *Eurycea cirrigera* (STP1715.1, one animal), and *Plethodon serratus* (STP1716.1, STP1717.1, STP1718.2, STP1718.4, three animals). General structural class was determined by manual analysis of GNPS library hits and outputs from Sirius 5.6.3 and CANOPUS for multiple nodes within a subnetwork.

A number of investigations of the skin microbiomes of frogs and other herptiles have identified specialized bacterial metabolites, for example, prodigiosin, violacein and volatile metabolites, such as antifungals against *Batrachochytrium* pathogens (56). In contrast, there appear to be no untargeted metabolomics studies of herptile gut microbiomes, and only two separate reports of specialized metabolites from cultured bacteria isolated from herptile guts. The latter publications document antibacterial activity of conditioned media filtrates from cultured gut bacteria and LCMS-based annotations of filtrate metabolites for a water monitor lizard (57) and a turtle (58). Subsequent antibacterial testing of commercially available metabolites annotated from the turtle-derived bacteria was also reported (59).

To our knowledge, antifungal basidiosins A-C are the only published specialized metabolites isolated and characterized from laboratory cultured *Basidiobolus* isolates (60). These structurally similar cyclic pentapeptides from *B. meristosporus* (isolate ARSEF 4516) contain both D and L amino acids. MK3990 (CAS # 136509-32-5) is another peptidic metabolite referenced in a 1991 Japanese patent application as an antibiotic produced by *B. meristosporus*, although no molecular structure is directly available. Notably, *Basidiobolus* genomes are enriched in specialized metabolite biosynthetic genes compared to other related fungi (53), and particularly in NRPS genes. Indeed, we have observed several peptidic metabolite signatures in untargeted mass spectrometry experiments and these metabolites are shared across strains of *Basidiobolus* isolated from different hosts, consistent with their genomic enrichment for NRPSs as a characteristic of the genus.

## MATERIALS AND METHODS

### Collection of animals and processing fecal samples

Animals included in this study were collected between 2014 - 2022. Collection information including host taxonomy (taxonomyTable S1) and geographic location of EPA Ecoregion III and IV levels. Details of animal collection and collection of fecal samples are provided in Walker et al. (2020), but briefly: Animals were collected into plastic bags with a small amount of field material (e.g., leaf litter). Each animal was given a unique collection number and the collection site was flagged with the same collection number. Animals were transported to processing sites (e.g., field labs) where they were removed from the field collection bags, placed in a new plastic bag, and surfaced washed with sterile dH_2_0 for approximately 1 min to remove debris and transient microbes. After washing an animal, a skin swab and a tail or toe clip was obtained, then the animal was placed in a moist chamber overnight. In the morning, fecal samples were collected with a sterile plastic scoopula. For animals collected from 2014 - 2018 fecal samples were placed into empty sterile microcentrifuge tubes, frozen and stored at -80°C until being thawed, diluted in 1 mL sterile molecular grade water, and processed using the protocol below. For animals sampled in 2022, fecal samples were placed into 1 mL of sterile molecular grade water, then vortexed for 20 sec and aliquoted as follows: 100 µl in 20% glycerol for culturing bacteria, 250 µl for DNA extractions, 250 µl for culturing *Basidiobolus*, and 400 µl for chemical analyses. Animals were then returned to their respective collection site after collection of fecal samples. A total of 33 different species were sampled from 16 frogs, 90 salamanders and 35 labsizards (Table S1).

### Culturing, microscopy, and imaging

Isolation of *Basidiobolus* was attempted for all 207 animals that produced fecal pellets in the 2022 field season. Canopy plates (61, 62) were prepared as follows: five ∼50 µL drops from the 250 µl fecal sample aliquot mentioned above were applied to the paper towel surface of the canopy plate. The plates were incubated with the paper towel (lid) surface side down with desk lamp illumination at ambient room temperature. Plates were monitored for *Basidiobolus* spore discharge over 2-5 days after which plates were autoclaved. Forcibly discharged blastoconidia with germinating hyphae were isolated from the PDA surface using sterile dissecting needles and stereoscope at ∼50X total magnification onto PDA plates.

Random selection of *Basidiobolus* isolates were grown in full strength PDA and Corn Meal Agar (CMA) for one week at 25°C. Fresh *Basidiobolus* cultures were used to prepare slide cultures (63) which were incubated at 25°C until the mycelia was observed on the cover slip. Coverslips were mounted on slides and stained with lactophenol cotton blue solution (Sigma) and observed under the light microscope. Microscope images were captured with a Leica DMC 4500 camera, using the Leica Application Suite v4.12.

### Amplicon sequencing and analysis - 16S rRNA and ITS rDNA markers

Target-gene data collection resulted in three data sets: 1) a broad sampling of 33 species of reptiles and amphibians that included 16 frogs, 90 salamanders, and 35 lizard individuals from the Midwestern, Southeastern, and Southwestern USA; 2) a selection of six *Basidiobolus* living strains isolated from different salamander species; and 3) a focused sampling of 60 Slimy Salamanders, which is a group of congeneric species of *Plethodon* that have only recently diverged and still hybridize, from 13 sites in the Southeastern USA (USATable S1).

DNA was extracted from fecal pellets using the Qiagen DNeasy 96 PowerSoil Pro kit and high-throughput sequencing completed on an Illumina MiSeq (2 x 250 bp paired-end) for the 16S rRNA V4 and ITS1 rDNA markers as in Walker et al. (2020). For bioinformatic analyses, primers of the 16S-V4 gene were removed from forward and reverse raw reads with Cutadapt v4.1 (64). Reads were filtered, dereplicated, trimmed (forward reads to 230 bp and reverse reads to 160 bp), and merged (min overlap of 100 bp) with R package DADA2 v1.24.0 (65). After inference of the Amplicon Sequence Variants (ASVs), chimeric sequences were removed, and taxonomy was assigned to each ASV using the naive Bayesian classifier method against the SILVA v.138 reference taxonomic database (66). ITS1 rDNA reads were first extracted from raw reads with ITSxpress v1.7.2 (67) and merged with BBMerge (68). Reads were filtered (min length of 50 bp) and ASVs were inferred with DADA2 v1.24.0 (65). After the removal of chimeric sequences, taxonomy was assigned to each ASV using the naive Bayesian classifier method against the UNITE+INSD fasta release v8.3 (69). For all datasets, the function *isContaminant* from the R package Decontam v1.16 (70) was used to identify and discard contaminant ASVs. Contaminants were identified using the option “method = frequency” which selects ASVs whose relative abundance varies inversely with sample DNA concentration. Decontamination was run at a range of probability threshold values (from 0.05 to 0.95, increasing by 0.10). This is the probability threshold below which the null-hypothesis (that an ASV is not a contaminant) should be rejected. For each dataset, we determined what percentage of sequences were removed from no-template control (NTC) libraries relative to sample libraries. A threshold value was selected if it was the last value at which more sequences were removed from NTC than sample libraries relative to the next value tested (given at least 10% removal of total NTC sequences). Both datasets were then rarified to a depth of 10,000 reads using the function *rrarefy* from the R package vegan v2.6-4 (71).

To understand intragenomic ITS rDNA allelic diversity, ITS rDNA amplicon sequencing was performed on six living isolates of *Basidiobolus*. Sequencing was performed as described in Walker et al. (2020), however, the bioinformatic analysis was completed as described above in DADA2.

The fungal ITS rDNA and 16S rRNA marker datasets were analyzed with R package Phyloseq v1.40.0 (72). The twenty most abundant fungal and bacterial genera were visualized in bar charts, highlighting differences among hosts and geography (states) (Fig. 2). Beta diversity of both datasets were inspected with a Principal Coordinate Analysis (PCoA) based on Bray-Curtis distances. A Betadisper analysis was performed to test for differences in multivariate dispersion between host groups. PERMANOVA tests (function adonis) were used to compare average microbiome assemblages among animal hosts, across geographic locations and the interaction between host and geography (R package Vegan v2.6.4) (71). Relative abundances of bacterial and fungal taxa among the different animal hosts were visualized with R package ampvis2 v2.7.34 (73).

### Sanger sequencing, phylogenetic reconstruction and ancestral state reconstruction of *Basidiobolus*

Genomic DNA was extracted using Extract-N-Amp method and the amplification of the ITS region from rDNA was performed with the primers ITS5 (5’-GGAAGTAAAAGTCGTAACAAGG-3’) and ITS4 (5’-TCCTCCGCTTATTGATATGC-3’) (74). The thermal cycling conditions were as follows: an initial denaturation step at 95°C for 2 min, followed by 30 cycles of 95°C for 30 sec, 55°C for 1 min and 72°C for 1 min, and a final extension step at 72°C for 5 min. PCR reactions (25 μL final volume) contained 2 μL of genomic DNA, 1.25 μL of each primer (10 μM each), 0.5 μL nucleotide mix (10 mM), 2.5 μL MgCl_2_ (25 mM), 5 μL GoTaq® Reaction Buffer (5X), 0.25 μL GoTaq® DNA Polymerase (5u/μL; Promega, Radnor, PA, USA), and 12.25 μL of molecular grade water. PCR products were cleaned with ExoSap-IT (Thermo Fisher Scientific, Waltham, MA, USA) and sequenced at the Center for Quantitative Life Sciences (Corvallis, Oregon, USA).

A total of 52 sequences from different species of *Basidiobolus* were downloaded from the NCBI GenBank database and aligned with the ITS sequences of *Basidiobolus* isolates. An ITS rDNA alignment was constructed using MAFFT v7 (75), and visualized and edited in Geneious Prime v2022.2 (https://www.geneious.com). A Maximum Likelihood phylogenetic tree was constructed with RAxML-HPC v8.0 (76) under GTR+GAMMA+I model with 1,000 bootstrap replicates (77). The final tree was visualized in FigTree v1.4.4 (http://tree.bio.ed.ac.uk/software/figtree/).

An Ancestral Character State Reconstruction (ACSR) analysis was performed on a selected group of *Basidiobolus* isolates to investigate the association of different hosts and geographic location (Ecoregion level III) at specific nodes. All ITS sequences used to construct the phylogenetic tree were clustered into Operational Taxonomic Units (OTUs) at 99% similarity using VSEARCH v2.22.1 (78). To maintain representation of hosts for each OTU, an ITS sequence was included from each *Basidiobolus* isolated from a unique host genus within each 99% OTU, resulting in seventy-three sequences. These sequences were aligned and a phylogenetic tree was constructed as previously described. Using FigTree, the phylogenetic tree was exported in nexus format for an ACSR analysis using the R Phytools package v1.2-0 (79). The results of ACSR analysis on host type and geographic location were displayed in two phylogenetic trees.

The phylogenetic tree constructed with the selected ITS sequences, was imported into Geneious v2023.0.4 (https://www.geneious.com), and a patristic distance matrix was calculated. The patristic distance matrix was used to test the effect of host groups, host genera and geographic location on the genetic change represented in the ACSR phylogenetic tree. A Betadisper analysis was used to test for differences in multivariate dispersion between host groups. PERMANOVA (function adonis) was used to test the effect on the phylogenetic change among hosts groups (salamanders, frogs, lizards), host genera, across geographic locations (Ecoregion III), and all interactions (R package Vegan v2.6.4) (71).

### *Basidiobolus* OTU – bacterial community correlations

A co-occurrence network analysis was performed on the complete dataset of frogs, lizards and salamanders with a frequency threshold of ASVs present in more than 20% of samples. After Clr transformation, the sparse inverse covariance estimation and model selection were implemented using the *spiec.easi* function in SPIEC-EASI with MB neighborhood selection (80). The *nlambda* was set for each model to obtain at least 0.49 or the closest possible value to the target stability threshold of 0.05. Data processing and networks construction were performed using R (version 4.2.2; R Core Team, 2022) in the packages phyloseq v1.42.0 (72), SpiecEasi v1.1.2 (80), igraph v1.3.5 (81) and microbiomeutilities v1.00.17 (82) and their dependencies. Network properties including clustering coefficient, edge density, connectivity and betweenness were calculated in igraph and cytoscape. The module/submodule detection and modularity analyses were performed using fast greedy modularity optimization as a function in igraph (83). Networks were visualized in Cytoscape v3.9.1 (84) using the edge-weighted spring embedded layout. We removed ASVs with <10 reads and 0.5% minimum abundance threshold to identify keystone bacterial or fungal ASVs in herptile gut microbiomes. The keystone microorganisms were identified by two parameters of within-module connectivity and among-module connectivity (46, 47).

Amplicon data from Walker et al. (2020) including the 16S rRNA and ITS1 rDNA were analyzed as 97% OTUs for 60 Slimy Salamander fecal samples. These data were utilized to explore the extent of correlation between bacterial taxa and *Basidiobolus* OTUs in a congeneric host (*Plethodon* spp.) (17). Rare bacterial and *Basidiobolus* OTUs (< 10 observations in dataset) were removed prior to downstream analyses. An indicator power analysis (44) was used to determine the ability of the 100 most abundant bacterial OTUs to predict the presence/absence of the most abundant *Basidiobolus* fungal OTUs (reads ≥ 10; n = 21 OTUs). The average indicator power value was calculated for each bacterial OTU. These values were then grouped by bacterial class to determine if there was taxonomic variation in the ability of bacterial OTUs to predict *Basidiobolus* occurrence. Total indicator power (TIP), the average ability of the members of an indicator assemblage to predict the occurrence of a target taxon, was calculated for each *Basidiobolus* OTU.

### Molecular network analyses

Ten different *Basidiobolus* isolates were grown over cellophane on potato dextrose agar for 21 days. Fungal mycelium was then collected and freeze dried before addition of HPLC grade MeOH (0.25 g of mycelium/mL). Suspensions were then sonicated for 30 min and left overnight. The extract was then filtered to remove mycelium and concentrated under reduced pressure. The extraction procedure and following analysis was also performed on a blank sample (empty vial) as a control. For tandem mass spectrometry analysis, fungal extracts were re-dissolved in LCMS grade MeOH (1 mg/mL) spiked with two internal standards (D-Ala2-odoamide (85), *m/z* 856.5474, 0.005 mg/mL; Tsn-Pc-832A (86), *m/z* 832.5404, 0.005 mg/mL). Full (0.5 mg/mL) and half (0.25 mg/mL) strength quality control samples containing six randomly chosen extracts were run at the beginning and end of the batch, and samples were run in random order, with a blank run every ten samples. For each chromatographic run, 3 µL of sample was injected on an Agilent 1260 infinity II LC coupled to a 6545 QToF MS. For the chromatographic separation, a reversed phase C18 porous core column (Kinetex C18, 50 x 2.1 mm, 2.6 µm particle size, 100 Å pore size, Phenomenex, Torrance, USA) was used. The mobile phase consisted of solvent A) H_2_O + 0.1 % formic acid (FA) and solvent B) acetonitrile (ACN) + 0.1 % FA, and the flow rate was 0.4 mL/min. After injection, the samples were eluted with a linear gradient from 0-0.5 min at 25 % B, 0.5-7 min 25-95 % B, 7-8 min 95 % B, followed by a 3.5 min washout phase at 100% B and a 5 min re-equilibration phase at 25 % B. The column compartment was maintained at 30 °C. Data dependent acquisition (DDA) of MS^2^ spectra was performed in positive mode. Electrospray ionization (ESI) parameters were set to a gas temperature of 325 °C, gas flow of 10 L/min, nebulizer 20 psi, sheath gas temperature of 375 °C, and a sheath gas flow of 12 L/min. The spray voltage was set to 600 V. MS scan range was set to *m/z* 100-3000 and the scan rate was 10 spectra/sec. Collision energy was set to a stepwise increase from 20 to 40 to 60 eV. MS^2^ scans were selected when precursor counts reach 1000 counts and spectra were excluded after six were collected. For MS^2^ data analysis, raw spectra were converted to to.mzML files using MSconvert (ProteoWizard). MS^1^ and MS^2^ feature extraction was performed using MZmine2.53. Each feature ID represents a m/z-retention time pair and has an associated MS^2^ spectrum and quantification across samples based on area under the chromatographic peak. The parameters used in MZmine2.53 are listed in inTable S2. The feature tandemblee tandemblee.csv and .mgf files were exported and uploaded to GNPS (gnps.ucsd.edu) (55) for feature-based molecular networking (FBMN) (54). For spectrum library matching and spectral networking, the minimum cosine score to define spectral similarity was set to 0.7. The precursor and fragment ion mass tolerances were set to 0.02 Da, minimum matched fragment ions to six. Molecular networks were visualized with Cytoscape 3.9.1 (84) and node information was enriched with the MS1 peak areas from the feature table.

Input Input.mgf files for Sirius 5.6.3 were exported from MzMine and opened in the Sirius 5.6.3 GUI (87). Jobs were run and filtered by an isotope pattern with an MS^2^ mass accuracy of 10 ppm. MS^2^ isotope scorer was ignored, and 10 candidates were stored for each mass feature. Possible ionizations included [M+H]^+^, [M+Na]^+^, and [M+K]^+^. ZODIAC (88) and CANOPUS (87, 89, 90) jobs were included with preset parameters. Masses greater than 850 Da were excluded.

## DATA AVAILABILITY

Raw ITS sequences from isolates, raw ITS and 16S amplicon sequence reads have been deposited in the NCBI/SRA database under the project accession PRJNA932855. Scripts and input data used for the analysis are available via a GitHub repository (https://github.com/herptilemicrobiomes/HerptileGutMicrobiome_2023). The link to the GNPS job is as follows: https://gnps.ucsd.edu/ProteoSAFe/status.jsp?task=ab180f58f4ee484683b7950461b22566.

## ACKNOWLEDGEMENTS

The authors would like to thank Aaron Ross, Clay Stalzer, Ian Trautman, Matt Brown, Matt Grisnik, Nathan Wilhite, Ori Bergman, Brandon Ison and Paul Super for support in field and/or lab activities. This work was supported by National Science Foundation grants EF-2125065, EF-2125066, EF-2125067 to D. Walker, J.W. Spatafora, J.E. Stajich & K.L. McPhail, and the Molecular Biosciences Program at MTSU. All research was conducted under IACUC22-3002, TWRA5107, TDEC2021-034, CHE901200, 14-SC00873, TWRA3886, AL2014044693068680, AL2016021753868680, 29-WJH-16-184, MS0722163.

## AUTHOR CONTRIBUTIONS

LVG, ASR, JS, JES, KLM, DMW drafted the manuscript, LVG handled major revisions, all authors contributed to data collection, analysis, and manuscript revisions.

